# EMT and MET: necessary or permissive for metastasis?

**DOI:** 10.1101/122051

**Authors:** Mohit Kumar Jolly, Kathryn E Ware, Shivee Gilja, Jason A Somarelli, Herbert Levine

**Affiliations:** Center for Theoretical Biological Physics, Rice University, Houston, TX 77005, USA; Duke Cancer Institute & Department of Medicine, Duke University Medical Center, Durham, NC 27708

**Keywords:** Epithelial-to-mesenchymal transition, mesenchymal-to-epithelial transition, metastasis, hybrid epithelial/mesenchymal, phenotypic plasticity

## Abstract

Epithelial-to-mesenchymal transition (EMT) and its reverse mesenchymal-to-epithelial transition (MET) have been often suggested to play crucial roles in metastatic dissemination of carcinomas. Recent studies have revealed that neither of these processes is binary. Instead, carcinoma cells often exhibit a spectrum of epithelial/mesenchymal phenotype(s). While epithelial-mesenchymal plasticity has been observed pre-clinically and clinically, whether any of these phenotypic transitions are indispensable for metastatic outgrowth remains an unanswered question. Here, we focus on epithelial-mesenchymal plasticity in metastatic dissemination and propose alternative mechanisms for successful dissemination and metastases beyond the traditional EMT-MET view. We highlight multiple hypotheses that can help reconcile conflicting observations, and outline the next set of key questions that can offer valuable insights into mechanisms of metastasis in multiple tumor models.

## Introduction

Epithelial-to-mesenchymal transition (EMT) is a cellular process loosely defined as a loss of the epithelial traits of tight cell-cell adhesion and apico-basal polarization and a gain of mesenchymal traits of motility and invasion (Savagner, 2015). The concept of EMT evolved from initial observations that embryonic and adult epithelial cells converted to migratory and invasive fibroblast-like cells when embedded in 3D collagen gels (Greenburg and Hay, 1982). Defined then as a ‘transformation’, EMT has since been well-studied in gastrulation, neural crest migration, heart development, branching morphogenesis, wound healing, fibrosis, and cancer metastasis. ‘Transformation’ has given way to ‘transition’ and more recently ‘plasticity’ to accurately represent its reversibility as well as its non-binary nature (Jolly et al., 2015a; Nieto et al., 2016). In the context of cancer, the proposition of EMT and MET driving the invasion-metastasis cascade (Thiery, 2002) has been pursued enthusiastically for over a decade (Hartwell et al., 2006; Jung et al., 2008; Mani et al., 2007; Ocaña et al., 2012; Onder et al., 2008; Spaderna et al., 2008; Stankic et al., 2013; Tsai et al., 2012; Yang et al., 2004), but recent studies have questioned the indispensability of these transitions in establishing metastasis (Fischer et al., 2015; Shamir et al., 2014; Somarelli et al., 2016a; Zheng et al., 2015). These results have stimulated provocative discussions on what steps are necessary and sufficient to establish macrometastases *in vivo*. Here, we attempt to reconcile some apparent contradictions, and highlight key unanswered questions that need to be addressed for a better understanding of the contribution of EMT and MET in metastasis in multiple tumor types.

### EMT and MET are not binary processes

A tacit assumption in the proposed role of EMT and MET during the metastasis-invasion cascade was that, similar to the distinct developmental lineages – epithelium and mesenchyme – carcinoma cells can attain either a fully epithelial or a fully mesenchymal state (Thiery, 2002). This assumption was supported by the labeling of phenotypes co-expressing canonical epithelial and mesenchymal markers as ‘metastable’, strongly suggesting that these observations were a snap-shot *en route* to full EMT/MET and thus could not reflect a stable state or an end point of a transition in itself (Lee et al., 2006). Only recently has the concept of a hybrid epithelial/ mesenchymal (E/M) state been revisited in cancer (Bronsert et al., 2014; Chao et al., 2011; Grosse-Wilde et al., 2015; Huang et al., 2013; Lecharpentier et al., 2011; McCart Reed et al., 2016; Naber et al., 2013; Sampson et al., 2014; Schliekelman et al., 2015; Strauss et al., 2011), and shown to be stable over multiple passages *in vitro* (Jolly et al., 2016). This revised understanding of cancer cell plasticity has been at least in part driven by computational modeling efforts of EMT/MET regulatory networks (Jia et al., 2015; Li et al., 2016; Lu et al., 2013; Zadran et al., 2014) that have triggered investigations of single–cell phenotypes in terms of their EMT status (Andriani et al., 2016; Grosse-Wilde et al., 2015).

In the context of wound healing and embryonic development, the intermediate state(s) of EMT has (have) been well-studied (Arnoux et al., 2005; Futterman et al., 2011; Johnen et al., 2012; Kuriyama et al., 2014; Leroy and Mostov, 2007; Micalizzi et al., 2010; Revenu and Gilmour, 2009; Shaw and Martin, 2016; Somarelli et al., 2013). The idea that EMT need not be an ‘all-or-none’ process (Nieto, 2013) has motivated a detailed dissection of five major sub-circuits involved in EMT - basement membrane remodeling, motility, cell-cell adhesion, apical constriction, and loss of apico-basal polarity. These sub-circuits are inter-connected and overlapping, but intriguingly, no single EMT-inducing transcription factors (EMT-TF) is involved in all of these sub-circuits, highlighting the complexity of cellular plasticity even in relatively simpler organisms such as sea urchin (Saunders and McClay, 2014). These observations enable envisioning EMT as a process in five-dimensional space, where induction of different EMT-TFs may alter the positioning of epithelial cells in different ways in this space. Thus, EMT progression is not a unidimensional linear process, but navigation through a rugged highly nonlinear landscape (Figure 1). Also, though assumed here as independent axes, these five aspects of EMT may affect one another too, thus compounding non-linearity of the process. Therefore, it may be more tricky to define EMT, but for practical purposes, we will consider here single-cell migration and/or invasion with loss of cell-cell adhesion as EMT, as was postulated initially (Cheung and Ewald, 2014). This loss of cell-cell adhesion typically co-occurs with a decrease in other epithelial traits such as loss of apico-basal polarity, and a concomitant increase in genes often expressed specifically in mesenchymal cells and tissues (Kalluri and Weinberg, 2009).

Moreover, the epigenetic reprogramming accompanying many of these key developmental events may rewire EMT regulatory networks differently in different tissue types, further amplifying the heterogeneity and context-dependence of EMT states. For instance, in breast cancer, basal cells exhibit bivalent chromatin states, with both activating and repressive marks for a key EMT-TF, ZEB1, but luminal cells only have repressive marks for ZEB1 (Chaffer et al., 2013). Thus, basal cells are already poised to display stronger EMT traits upon exposure to EMT-inducing cytokines such as TGFβ, as compared to luminal cells. Similarly, GRHL2 – a transcription factor that can inhibit EMT (Varma et al., 2012; Walentin et al., 2015) – can be methylated in sarcomas as compared to carcinoma (Somarelli et al., 2016b), thus rewiring the circuit regulating EMT in sarcomas.

**Figure 1.**
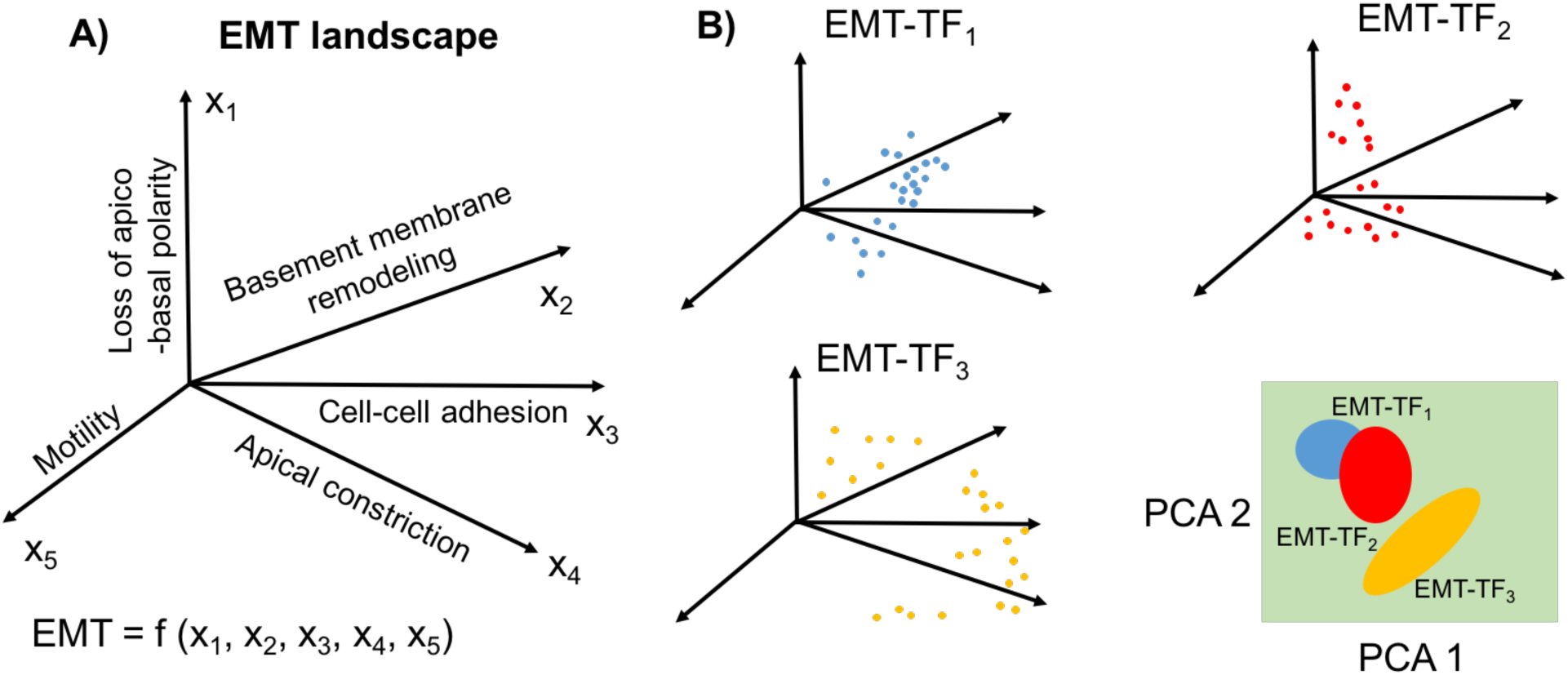
Representing EMT as a multi-dimensional non-linear process. **A)** EMT landscape may contain multiple axes (*x*_1_-*x*_5_). **B)** Induction of EMT by different EMT-TFs may drive epithelial cells into different regions on this multi-dimensional landscape (shown by different colored dots). There may be some overlap in the effect of more than one EMT-TFs in regulation of one or more of these axes contributing to EMT, as can be realized by projecting this multi-dimensional space into two principal component axes (PCA).

Therefore, with such ubiquitous tissue- or even subtype-specific complexity and heterogeneity being revealed, binning carcinoma phenotypes into either fully epithelial or fully mesenchymal, and dismissing all hybrid phenotypes as ‘metastable’, can only hamper a better understanding of both the nuances of EMT and MET, and how these processes may impinge on metastasis.

### Role of EMT-TFs in metastasis: necessary or permissive?

In the context of cancer, Snail (SNAI1) was identified as the first EMT-TF that directly repressed transcription of the epithelial cell-cell adhesion molecule, E-cadherin. Overexpression of Snail in MDCK and many carcinoma cell lines led to loss of cell-cell adhesion mediated by E-cadherin, transformed the morphology of cells from epithelial to spindle-liked mesenchymal, and enhanced their migratory and invasive traits *in vitro* (Batlle et al., 2000; Cano et al., 2000). Further work revealed a similar, but less potent role of another EMT-TF Slug (SNAI2, a member of *Snail* family) both *in vitro* and *in vivo* (Bolós et al., 2003; Hajra et al., 2002; Ware et al., 2016). Snail was also shown to induce the expression of mesenchymal markers fibronectin and Zeb1 (Guaita et al., 2002), the latter of which is another EMT-TF that can promote tumor invasiveness *in vitro* and is correlated with tumor cell differentiation *in vivo* (Aigner et al., 2007; Spaderna et al., 2008). Later, Twist was identified as yet another EMT-TF that inhibited E-cadherin as well as regulated other components of EMT in MDCK, mammary epithelial cells (Yang et al., 2004) and breast cancer cell lines (Vesuna et al., 2008). Silencing Twist in 4T1 cells abrogated the number of lung metastases significantly, however, did not completely abolish them (Yang et al., 2004), still leaving open the possibility that Twist, and potentially other EMT-TFs, may act more as accelerators rather than drivers of metastasis (Figure 2A). These above mentioned studies confirmed that the EMT-TFs that governed developmental EMT also contributed to one or more aspects of EMT in carcinomas *in vitro*, a claim that was substantiated by *in vivo* negative correlation between these EMT-TFs and E-cadherin expression. Thus, these studies led to a conceptual framework suggesting that aberrant activation of one or more EMT-TFs (resulting from many potential scenarios such as hypoxia, secreted EMT-inducing cytokines from the stroma, e.g. TGF-β, altered degradation rate of EMT-TFs etc.) was a necessary and sufficient condition for metastasis.

**Figure 2.**
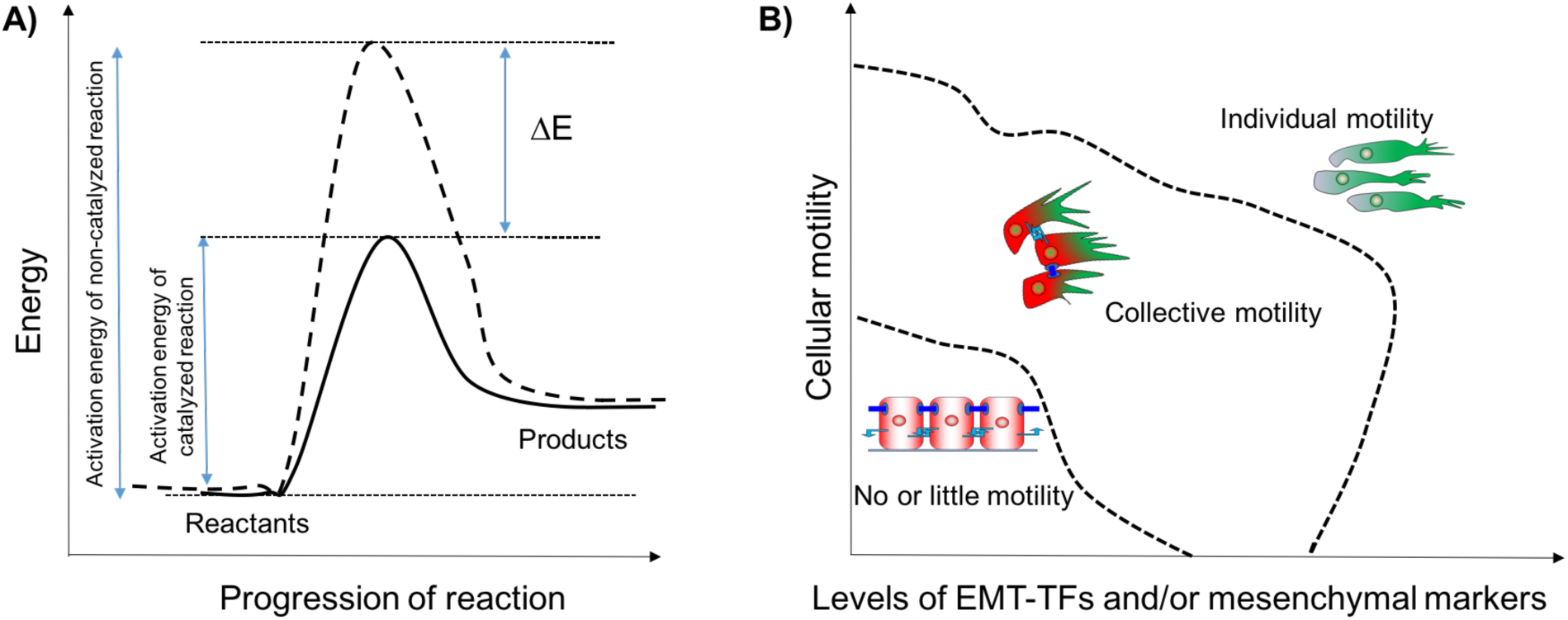
Role of EMT-TFs. **A)** EMT-TFs can act as catalysts of cellular plasticity. A catalyst reduces the activation energy (by an amount of ΔE) required for the progression of a reaction. **B)** Phase diagram showing different types of motility that can be possible at varying levels of EMT-TFs and/or mesenchymal markers, and cellular motility. Dotted lines represent phase separations.

This framework was recently challenged by two lineage-tracing studies in mouse models of pancreatic and breast cancer. Zheng *et al*. genetically knocked out either Twist or Snail in spontaneous pancreatic ductal adenocarcinoma (PDAC) models, but the incidence of metastasis was not altered significantly (Zheng et al., 2015). Multiple alternative interpretations have been proposed for this observation – (a) knockdown of one EMT-TF need not be sufficient to ablate EMT completely, and compensatory EMT-inducing pathways may be present (Li and Kang, 2016), (b) the marker used for lineage-tracing of cells undergoing EMT in this study – α smooth muscle actin (αSMA) – is rarely induced spontaneously upon activation of EMT in this particular mouse model (Pattabiraman and Weinberg, 2017). The other study (Fischer et al., 2015) focused on spontaneous breast-to-lung metastasis mouse models and used Fibroblast-specific protein1 (Fsp1) as a lineage-tracing marker of cells undergoing an EMT. They found many Fsp1-negative cells metastasizing to lung, suggesting that not even a transient activation of EMT was essential for metastasis. The specificity and sensitivity of Fsp1 to mark cells undergoing EMT and/or fibroblasts may be called into question (De Chiara and Crean, 2016; Pattabiraman and Weinberg, 2017). But this study also demonstrated that overexpression of miR-200 suppressed multiple levels of many EMT-TFs including Zeb1 yet did not affect lung metastasis (Zheng et al., 2015), thus providing a stronger argument for alternative mechanisms of dissemination.

In contrast, knockdown of Zeb1 in HCT116 and SW480 cells has been shown to abolish lung metastases after intrasplenic or intravenous injection in nude mice (Spaderna et al., 2008). Similarly, deletion of TWIST1 drastically inhibited lung metastasis of 4T1 cells implanted in mice mammary gland (Yang et al., 2004), emphasizing a causal role of EMT-TFs in metastasis. Technically speaking, these studies were done with different approaches than in spontaneous metastasis genetically engineered mouse models (GEMM) discussed earlier. Nevertheless, it is likely that metastasis for all carcinoma cells need not require an overt upregulation of various EMT markers to gain migratory and invasive traits. For instance, in tumor organoids, breast cancer cells can invade the extracellular matrix (ECM) by three modes - collective invasion, mesenchymal invasion, and amoeboid invasion. In this model system, only cells undergoing mesenchymal invasion utilize an EMT-like program (Nguyen-Ngoc et al., 2012). Conversely, collectively invading cells do not typically express Vimentin or Twist1 and maintain E-cadherin mediated contacts with follower cells. Rather than undergoing an EMT, the cells undergoing collective invasion appear to undergo a transition toward a more basal-like phenotype, expressing K14 and p63 (Cheung et al., 2013). Put together, it still remains a possibility that the traits needed for successful metastasis can be gained by altering cellular adhesion and invasion through pathways that do not necessitate supra-physiological or aberrant overexpression of one or more EMT-TFs identified so far (Figure 2B). In other words, morphological changes associated with EMT can occur without an overt upregulation of any mesenchymal markers (Cheung and Ewald, 2014). Further, an overt or a complete EMT may not be as efficient for metastasis as retention of some molecular and/or morphological epithelial traits (Jolly et al., 2015a; Shamir et al., 2014).

### Has a full EMT ever been seen *in vivo*?

Recent progress in considering EMT as more of a spectrum of phenotypes instead of a binary process has driven an emerging notion that unlike during development, in which terminally differentiated epithelial and mesenchymal states exist, carcinoma cells might undergo more partial transitions to an incomplete mesenchymal phenotype (Lambert et al., 2017; Nieto et al., 2016). This notion is supported by observations that induction of a fully mesenchymal state through overexpression of an EMT-TF may lead to a loss of tumor-initiation potential and thus the ability to colonize (Celià-Terrassa et al., 2012; Ocaña et al., 2012; Ruscetti et al., 2015; Tsai et al., 2012). Earlier studies based on similar overexpression of EMT-TFs proposed an increase in tumor-initiation potential (Mani et al., 2008). Reconciling these contradictions, recent studies that categorized cells into E (epithelial), M (mesenchymal) and hybrid E/M, instead of just E and M have proposed that tumor-initiation potential might be maximum when cells are in a hybrid E/M state (Grosse-Wilde et al., 2015; Jolly et al., 2014; Ombrato and Malanchi, 2014; Ruscetti et al., 2015). Such hybrid E/M cells co-expressing various epithelial and mesenchymal markers have been observed in cell lines of breast, ovarian, lung, and renal cell carcinomas (Andriani et al., 2016; Grosse-Wilde et al., 2015; Huang et al., 2013; Sampson et al., 2014; Schliekelman et al., 2015), in mouse models of prostate cancer and PDAC (Rhim et al., 2013; Ruscetti et al., 2015), primary breast and ovarian cancer tissue (Strauss et al., 2011; Yu et al., 2013), bloodstream of breast, lung, and prostate cancer patients (Armstrong et al., 2011; Lecharpentier et al., 2011; Yu et al., 2013), and metastatic brain tumors (Jeevan et al., 2016). More importantly, triple negative breast cancer patients had a significantly higher number of such hybrid E/M cells as compared to other subtypes, suggesting a correlation between a hybrid E/M phenotype and tumor aggressiveness (Yu et al., 2013).

Although it is likely that many carcinomas undergo only a partial transition, some cancers reflect a more complete phenotypic transition based on typical morphological and molecular readouts. For example, Beerling *et al*. identified a rare population of E-cad^lo^ cells that underwent spontaneous full EMT without any experimental induction of EMT-TFs, and converted to being epithelial upon reaching the metastatic site (Beerling et al., 2016). Another model system that tends to exhibit a (complete) EMT is the Dunning model of prostate cancer that was derived in 1961 from a spontaneous prostate adenocarcinoma in a Copenhagen rat (Dunning, 1963; Issacs et al., 1978). The DT cell line established from this model expresses numerous epithelial biomarkers, including E-cadherin, claudin 4, and pan-cytokeratin (Oltean et al., 2008), possesses a cobblestone-like appearance (Oltean et al., 2006; Somarelli et al., 2013, 2016a), and, when implanted back into syngeneic rats, produces an extremely slow-growing, indolent tumor (Presnell et al., 1998). Serial passage in castrated rats of this tumor led to a diverse family tree of increasingly aggressive tumors and derivative cell lines (Issacs et al., 1982; Smolev et al., 1977; Tennant et al., 2000). One of these cell lines, derived from an anaplastic, highly aggressive variant, led to the development of the anaplastic tumor 3 (AT3) cell line. Compared to pre-EMT DT cells, AT3 cells exhibit a post-EMT phenotype (Oltean et al., 2008, 2006; Somarelli et al., 2013), with spindle-like morphology, low cell-cell attachment, enhanced invasion (Schaeffer et al., 2014), and metastatic capacity (Oltean et al., 2006). Consistent with these observations, microarray analysis of DT and AT3 cells revealed distinct epithelial and mesenchymal biomarker expression, with robust expression of multiple epithelial markers in the DTs and mesenchymal markers in the AT3s (Oltean et al., 2008). These analyses suggest that *in vivo* serial passage under androgen-deprived conditions induce a phenotypic transition consistent with EMT in the AT3 line. Thus, AT3 cells tend to reflect ‘epigenetically fixed’ EMT, similar to earlier where three unique phenotypic states were reported - epithelial, metastable E/M, and an ‘epigenetically fixed’ mesenchymal (Thomson et al., 2011). Yet, unlike the findings discussed above in which a complete EMT reduces the metastatic capacity of the cells, AT3 cells are highly metastatic and remain in a ‘fixed’ mesenchymal state during metastatic colonization ( Somarelli et al.,2016a).

Clinically, EMT has been suggested to play a role is in the case of carcinosarcomas - rare cancers comprised of both carcinomatous and sarcomatous elements. Genetic analyses supports a clonal origin of both epithelial and stromal elements within these tumors (Somarelli et al., 2015). Interestingly, cells expressing markers and/or morphological features of an intermediate or hybrid epithelial/mesenchymal state have been observed (Bittermann et al., 1990; DeLong et al., 1993; Haraguchi et al., 1999; Paniz Mondolfi et al., 2013), suggesting that the mesenchymal component is derived via EMT from the carcinomatous component. While it remains to be conclusively tested whether carcinosarcomas represent tumors in which a portion of the cells underwent EMT, the majority of data suggest that, in most cases, the mesenchymal element is likely derived from a carcinoma (Somarelli et al., 2015).

Similar to carcinosarcomas, in which tumors exhibit admixture of two phenotypes, prostate tumors with areas of adenocarcinoma and neuroendocrine prostate cancer (NEPC) have also been observed. Both the adenocarcinomatous and NEPC phenotypes share common mutations, suggesting a common cell of origin (Beltran et al., 2011; Hansel et al., 2011; Tan et al., 2014). Likewise, a longitudinal analysis of patients with adenocarcinoma that progress to NEPC indicated that NEPC results from clonal evolution of an original adenocarcinoma through phenotypic plasticity (Beltran et al., 2016). Further lineage-tracing studies support this finding, with combined genetic loss of Pten/Rb1/Trp53 inducing an NEPC-like transition by upregulating stemness factor Sox2, and epigenetic remodeling protein Ezh2 (Ku et al., 2017; Mu et al., 2017). While not a classic example of EMT, NEPC-like tumors represent similar phenotypic plasticity, and some players implicated in EMT such as Snail have been also reported in the context of NEPC-like tumors and neuroendocrine differentiation (McKeithen et al., 2011).

Taken together, although induction of at least a partial EMT at the invasive edges in primary xenografts has been observed *in vivo* (Bonnomet et al., 2012; Klymkowsky and Savagner, 2009), a careful investigation of partial vs. full EMT needs to be conducted *in vivo* to dissect the contributions of these phenotypic transitions to invasion, dissemination and metastasis. It is also likely that each tumor’s requirements for EMT/MET are slightly different depending on the original cell of origin (e.g. basal vs. luminal), its unique mutation profile (e.g. p53 loss), and its epigenetics (e.g. bivalent vs. monovalent chromatin). A more sophisticated understanding of the hybrid E/M phenotype and its molecular underpinnings will surely help to further elucidate the context-dependent requirements for plasticity during various stages of the metastatic cascade.

### Cohesive cell migration and EMT: mutually exclusive migration modes?

Many recent reports have suggested alternative mechanisms for the escape of carcinoma cells, besides the single-cell dissemination as enabled by EMT. Specifically, collectively invading cells have been shown to migrate through the extracellular matrix (ECM) with intact cell-cell junctions (Clark and Vignjevic, 2015; Friedl et al., 2012). Collective invasion need not always exhibit significant changes in canonical epithelial and mesenchymal markers (Cheung et al., 2013; Shamir et al., 2014), but cells at leading edge of these cohorts may express certain EMT traits (Westcott et al., 2015). A three-dimensional reconstruction of serial section samples of many tumors has suggested that cell clusters are the predominant agents of invasion and that singlecell dissemination is extremely rare (Bronsert et al., 2014). Some of these collectively invading cohorts – referred as ‘tumor buds’ – displayed loss of cell polarity, reduced total levels and membrane localization of E-cadherin, and increased nuclear ZEB1. However, because these cells were not spindle-shaped and maintained E-cadherin levels at least partially, they were labelled as a hybrid E/M phenotype, instead of a full EMT (Bronsert et al., 2014; Grigore et al., 2016). It is expected that collectively invading strands and tumor buds are precursors of clusters of CTCs, also called as tumor emboli, as observed in patients with invasive melanoma, lung cancer, inflammatory breast cancer, and clear cell renal cancer (Hou et al., 2012; Jolly et al., 2015a; Kats-Ugurlu et al., 2009; Ye et al., 2010), thereby suggesting that the clusters of tumor cells retaining some of their epithelial traits can complete the metastasis-invasion cascade and give rise to polyclonal metastatic colonies (Cheung et al., 2016). However, whether the clusters need upregulation of any mesenchymal markers still remains to be investigated extensively.

These clusters of CTCs, although much less prevalent than individually migrating CTCs, can act as primary ‘villains’ of metastasis by forming 50-times more tumors as compared to individual CTCs (Aceto et al., 2014). Besides, clusters may be more efficient in resisting cell death during circulation and associate with significantly worse outcome in patients (Cheung and Ewald, 2016). Inhibiting players that mediate cell-cell adhesion directly or indirectly in these clusters such as plakoglobin or keratin 14 (K14) compromised their metastatic potential (Aceto et al., 2014; Cheung et al., 2016). These results are reminiscent of the essential role of E-cadherin in forming tumor emboli and distant metastasis in inflammatory breast cancer (Tomlinson et al., 2001) – a highly aggressive cancer that predominantly metastasizes via clusters (Kleer et al., 2001). Thus, retention of cell-cell adhesion as an epithelial trait may actually be crucial to successful metastasis in many aggressive cancers.

Activation of an EMT program – either fully or partially – at the invasive edge can alter the ability of primary tumor cells to intravasate and disseminate as individual CTCs (Bonnomet et al., 2012; Roth et al., 2016), and CTCs can display a dynamic spectrum of EMT phenotypes (Yu et al., 2013). But, any causal role of EMT-TFs, and by extension, of a partial or full EMT in mediating CTC cluster formation still remains to be thoroughly investigated. This issue is convoluted by observations that CTC clusters may contain platelets that are known to secrete TGF-β (Aceto et al., 2014), a potent mediator of EMT. Recently developed technologies to isolate CTC clusters, such as Cluster-Chip, may be critical in this endeavor (Sarioglu et al., 2015).

### Is MET required for metastasis?

While many studies have focused on the importance of EMT during metastasis (Tsai and Yang, 2013), it has also been hypothesized that cells transition back to an epithelial state through MET to form macrometastases (Thiery, 2002). This hypothesis is based upon the observation that many metastases express epithelial markers (Christiansen and Rajasekaran, 2006).

For example, Chao *et al*. examined E-cadherin expression in primary breast tumors and matched metastases and found 62% of cases had increased E-cadherin at the metastatic site compared to the primary tumor (Chao et al., 2010). Although metastatic tumors commonly display an epithelial phenotype, it has also long been known that undifferentiated/ mesenchymal metastases also occur in cancer patients. Even in a single patient there is heterogeneity in the phenotypic status of multiple metastases (Spremulli and Dexter, 1983). These observations lead us to inquire about the requirement of MET for metastasis. Do some disseminated tumor cells not require MET to colonize secondary sites? Or do colonized tumor cells retain a high level of phenotypic plasticity, thereby priming them for multiple rounds of MET and EMT subsequent to metastatic seeding?

Brabletz postulated two types of metastatic progression - one based on phenotypic plasticity and the other plasticity-independent. Metastatic progression that is based on phenotypic plasticity would require MET in order to colonize secondary sites. On the other hand, tumor cells can acquire genetic alterations that give all the necessary traits for dissemination and metastatic seeding in one go and do not require MET (Brabletz, 2012). *In vivo* experimental evidence for these two models of metastatic progression was demonstrated using lethal reporters of MET that kill all the cells undergoing MET. These reporters revealed the existence of both MET-dependent and MET-independent paths to metastatic progression – an MET-dependent path in carcinosarcomas, whereas an MET-independent path in prostate cancer (Somarelli et al., 2016a). It is likely that EMT-TFs and microRNA families that maintain an epithelial phenotype (Bracken et al., 2008; Burk et al., 2008; Lu et al., 2013) regulate MET-dependent metastatic mechanisms. Indeed, it was recently shown in a spontaneous squamous cell carcinoma model that Twist1 activation promoted EMT and CTCs. However, turning off Twist1 at distant sites allowed MET and was essential for disseminated tumor cells to proliferate and form macrometastases (Tsai et al., 2012), reminiscent of observations that EMT typically arrests the cell cycle (Vega et al., 2004).

Mechanisms underlying MET-independent metastasis still remain elusive. One hypothesis is based on recent observations that cells that fail to undergo cell cycle arrest upon induction of EMT accumulate genomic instability (Comaills et al., 2016). Therefore, the cells metastasizing independent of MET may be genomically unstable. This instability may serve to enrich for the rare subset of cells that are likely to lead to dedifferentiated and highly metastatic tumors that are cross-resistant to next-line therapies (Creighton et al., 2009; Sun et al., 2012). Therefore, therapies used to treat cancer cells may also select for genetic alterations that allow for both the maintenance of an EMT and sustained uncontrolled proliferation, thus potentially obviating the need for MET.

An alternative explanation of the results presented above is that cells might undergo a partial MET, which reporters could miss capturing, just as many reporter systems may be less sensitive in capturing a partial EMT (Li and Kang, 2016; Pattabiraman and Weinberg, 2017). In partial MET, cells are likely to retain their mesenchymal traits and gain their proliferative ability without the acquisition of any genetic alterations. In a study comparing primary and metastatic tissue from breast and prostate cancer, E-cadherin was found at the cellular membranes more often in metastases than in primary tumors. However, metastases also retained mesenchymal markers Vimentin and Fsp1 (Chao et al., 2012). This study suggests that some metastases may maintain a high amount of phenotypic plasticity and are primed to switch between states as selection occurs during growth or by treatment. Thus, it is not necessarily the phenotype that favors metastasis, but the acquisition of the suite of traits needed to metastasize.

A central question that remains unanswered is whether partial EMT is the same as partial MET in its phenotypic consequence. Most phenotypic studies have been done in carcinomas, which are derived from epithelial cells. As discussed above, these cells likely retain intrinsic epithelial phenotype and acquire migratory and invasive traits, leading to a partial EMT that can promote tumor dissemination (Jolly et al., 2015a). Yet, since these cells are still epithelial in origin they are probably often less likely to undergo a complete epigenetic reprogramming similar to that in mesenchymal tissues. Thus, it is not surprising that many carcinoma cells revert to an epithelial-like state when arriving to an epithelial environment to form metastases. It is crucial that these cells are able to reactivate the cell cycle to proliferate and colonize; if the cells become fixed in a mesenchymal-like phenotype and break the connection between the epithelial phenotype and cell cycle activation, either by mutation or epigenetic reprogramming, their metastatic potential might be severely compromised (Figure 3).

**Figure 3.**
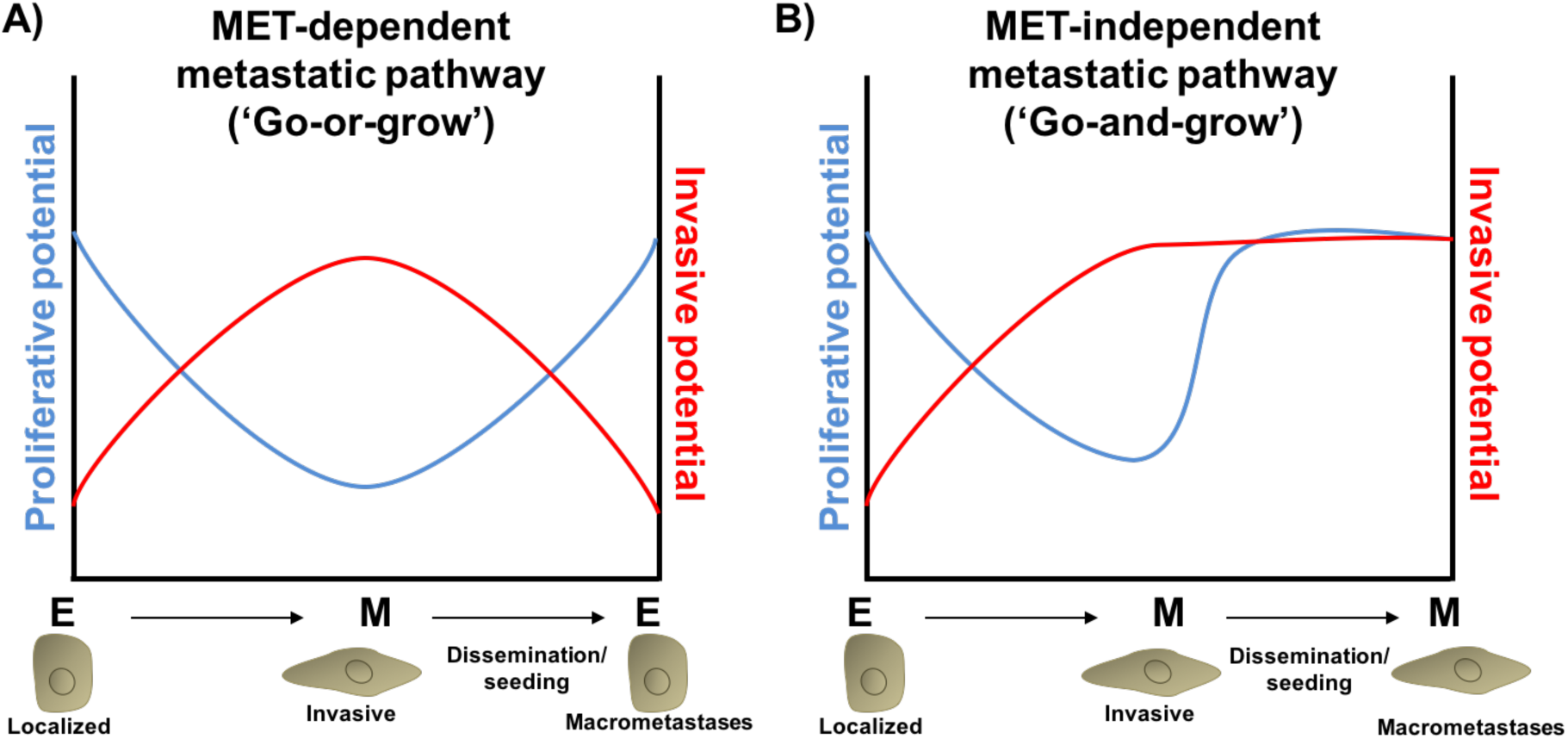
Plasticity-dependent and –independent pathways to metastasis. **A)** In MET-dependent metastasis, post-EMT-like cancer cells upregulate invasive programs that facilitate dissemination and seeding. The invasive program comes at a cost; EMT induction leads to downregulation of proliferative potential. Re-establishment of an epithelial-like phenotype via MET at the metastatic site awakens the proliferative potential necessary for formation of macrometastases. **B)** In MET-independent metastasis, therapy, epigenetic reprogramming, acquisition of novel mutations, or other mechanisms induce a post-EMT state that becomes fixed in a proliferation^*high*^/invasion^*high*^ phenotype. Cells metastasizing via an MET-independent pathway may be more aggressive, stem-like, chemo-refractory, and more likely to seed and re-seed further metastases.

Interestingly, sarcomas provide a unique perspective on the need for MET during metastasis. Sarcomas are cancers of a mesenchymal lineage. These cancers are highly aggressive and metastatic, and upregulation of mesenchymal biomarkers is observed in metastases compared to primary tumors (Shen et al., 2011; Wiles et al., 2013), suggesting that these tumors metastasize via an MET-independent route. It is possible that sarcoma cells are primed for enhanced metastatic capacity because of their mesenchymal lineage and that the acquisition of growth advantages during cancer initiation enables these cancers to metastasize readily via an MET-independent route. Clinically, sarcomas occur in younger patients and have a shorter overall survival compared to carcinomas (Siegel et al., 2017), suggesting that the rate-limiting step may be tumor initiation for metastasis. Conversely, in carcinomas, sustained cell growth is commonly coupled to MET during the formation macrometastases. In this scenario, induction of a MET might be the rate-limiting step in metastasis.

### Role of the microenvironment

Phenotypic plasticity can be influenced by tumor microenvironment; for instance, upregulation of hypoxia (Sun et al., 2009) and soluble factors released by macrophages and other infiltrating immune cells (Huang and Du, 2008; Toh et al., 2011) lead to upregulation of EMT-TFs and an induction of EMT. The importance of the microenvironment in driving a metastatic phenotype is underscored by the presence of ‘tumor microenvironment of metastasis’ (TMEM) – spatial proximity of an invasive cancer cell, a macrophage, and an endothelial cell, as identified by intravital imaging – being highly correlated with metastasis (Robinson et al., 2009). Therefore, normalizing the tumor microenvironment has been suggested to improve patient outcomes (Jain, 2013).

Dynamics of the microenvironment can enable a passive shedding of cancer cells into circulation. This mode would be instead of a postulated active crawling or migration of cancer cells towards any nutrient or chemokine gradient and cleavage of extracellular matrix (ECM) by secreting proteases (Bockhorn et al., 2007) For instance, blood vessels have been proposed to engulf clusters of cancer cells thus obviating the need for EMT (Fang et al., 2015). These clusters may avoid cell death in circulation by cell-cell contact mediated survival signals (Shen and Kramer, 2004), and may already be enriched for players such as Jag1 (Cheung et al., 2016) that can help them evade multiple therapies (Boareto et al., 2016; Li et al., 2014; Shen et al., 2015; Simões et al., 2015) and colonize successfully (Sethi et al., 2011). Not surprisingly, Jag1 is enriched in aggressive cancers such as basal-like breast cancer (BLBC) (Reedijk et al., 2008) and can contribute to the abnormal vasculature typically observed in cancers (Benedito et al., 2009; Boareto et al., 2015a). Moreover, Fringe, a glycosyltransferase that inhibits the binding of Notch to Jag1 (Boareto et al., 2015b; Jolly et al., 2015b), is lost BLBC (Zhang et al., 2014).

Therefore, it is not inconceivable that tumor cell dissemination – particularly cluster-based dissemination – is a passive process where cells that can navigate the fitness bottlenecks from an evolutionary standpoint eventually form metastases (Amend et al., 2016). Both genetic and non-genetic heterogeneity may be crucial or even synergistic in conferring a rare subpopulation of cells with high adaptability or plasticity that lets them transit the entire invasion-metastasis cascade. Such plasticity may coincide with co-expression of many epithelial and mesenchymal markers.

## Conclusion

Single-cell dissemination as enabled by EMT followed by a MET has been considered to be a hallmark of metastasis. However, alternative modes of dissemination such as collective or cluster-based migration and invasion can exist where cells need not shed cell-cell adhesion completely, and may not even exhibit an overt upregulation of mesenchymal markers, while having gained the traits of adhesion and invasion. Furthermore, migrating cells may lead to metastatic outgrowth without undergoing MET. Therefore, EMT and MET may not be playing a necessary role, but more of permissive and potentially catalytic roles, in metastatic dissemination of some carcinomas.

